# Tsunahiki task: A newly developed group-based operant task for mice

**DOI:** 10.64898/2026.01.09.697521

**Authors:** Mariko Nakata, Rino Iwabuchi, Tomoaki Murakawa, Shinnosuke Dezawa, Kotarou Hattori, Yui Kohno, Tsuyoshi Setogawa

**Affiliations:** Laboratory of Behavioral Neuroendocrinology, University of Tsukuba, Ibaraki 305-0006, Japan; Institute of Human Sciences, University of Tsukuba, Tsukuba, Ibaraki 305-0006, Japan; Human Informatics and Interaction Research Institute, National Institute of Advanced Industrial Science and Technology, Ibaraki 305-8566, Japan; Graduate School of Comprehensive Human Sciences, University of Tsukuba, Tsukuba, Ibaraki 305-0006, Japan; Faculty of Rehabilitation, R professional University of Rehabilitation, Ibaraki 300-0032 Japan; System Emotional Science, Faculty of Medicine, University of Toyama, Toyama 930-0194, Japan; Research Center for Idling Brain Science, University of Toyama, Toyama 930-0194, Japan

## Abstract

A group of social animals, including humans, distributes work and rewards and establishes inter-individual relationships through shared experiences of working together. Understanding the neural mechanisms underlying group work is essential for elucidating how social relationships are formed. Thus, a behavioral paradigm for studying group work in laboratory rodents is required. Here, we developed a Tsunahiki task, a novel group-based operant task for mice. In this task, three mice jointly pulled three ropes, and once all ropes were pulled out, all members gained access to a reward area, regardless of who performed the work. The mice acquired the task within a few days. Importantly, repeated experience with the Tsunahiki task led to a shift in workload toward subordinate individuals and induced rank consolidation. These findings suggest that group work induces consolidation of intra-group disparities based on the dominance hierarchy. The Tsunahiki task provides a useful framework for investigating the neurobiological mechanisms underlying collaborative work, group formation, and social inequality in rodents.

**Teaser:** A group-based operant task for mice was newly developed, in which hierarchy affected work and reward distributions.

## Introduction

We often work in groups. Working together enables us to accomplish highly advanced and complex tasks or reduce the burden by sharing workloads. The distribution of work and rewards in group work has always been of major interest in human society. They are usually determined through social interactions at the moment, and some individuals may shoulder more work or receive greater rewards. Numerous studies have aimed to reveal the underlying mechanisms of these complicated inter-individual interactions during group work and the influence of group work on social relationships. For example, in humans, the experience of working together promotes group formation (*1, 2*). Moreover, the establishment of a social hierarchy within a group through long-term repetition of group work has been reported in small-scale societies of forager-horticulturalists and nomadic pastoralists (*3*) and university students in the US (*4*).

Social interaction and establishment of relationships have been investigated in mice, a social species living in groups. Mice quickly establish hierarchical social relationships, where dominants acquire more resources, including food and mates, compared to subordinates (*5*) when they are housed with new members. Recently, experimental paradigms that require a group of mice to perform tasks have been developed. The IntelliCage® enables us to observe social behaviors over extended periods in the home cage of relatively large groups of 8 to 16 mice. During the observation, researchers can assign each mouse a learning task regarding the location of the water bottle (*6, 7*). In contrast, behavioral tasks originally developed for a single mouse have also been applied to multiple mice for simultaneous performance. Lara-Vasquez et al. (*8*) used a group of four mice to run T-maze simultaneously after each mouse learned a different “correct” arm. They demonstrated that individuals with a higher social rank within the group were influenced by other individuals compared to subordinates. However, in these behavioral tasks, even when tests are conducted in groups, learning occurs individually, with workload and rewards set for each subject. Moreover, although there are cooperative tasks that require two animals to do something simultaneously, such as the loose-string task (*9, 10*), few studies have been conducted in groups of three animals or more.

Here, we developed a group-based operant task to investigate group formation through task execution in mice (Fig. 1A). The task was conducted using a rectangular experimental arena divided into two areas: a start area and a reward area with a transparent guillotine door. At the beginning of the task trial, three mice were released from the start box, which was placed at the edge of the start area. They pull on ropes stuck through holes in the door. When they pulled out all three ropes within 20 minutes, the guillotine door was opened, and the mice could explore the reward area, where reward pellets were placed, for 5 minutes. All animals could enter the reward area, regardless of which individual pulled the ropes. Similarly, pellet acquisition was independent of individual task performance. We named this task the “Tsunahiki task.” *Tsunahiki* means rope-pulling or tug-of-war in Japanese.

**Fig. 1.**
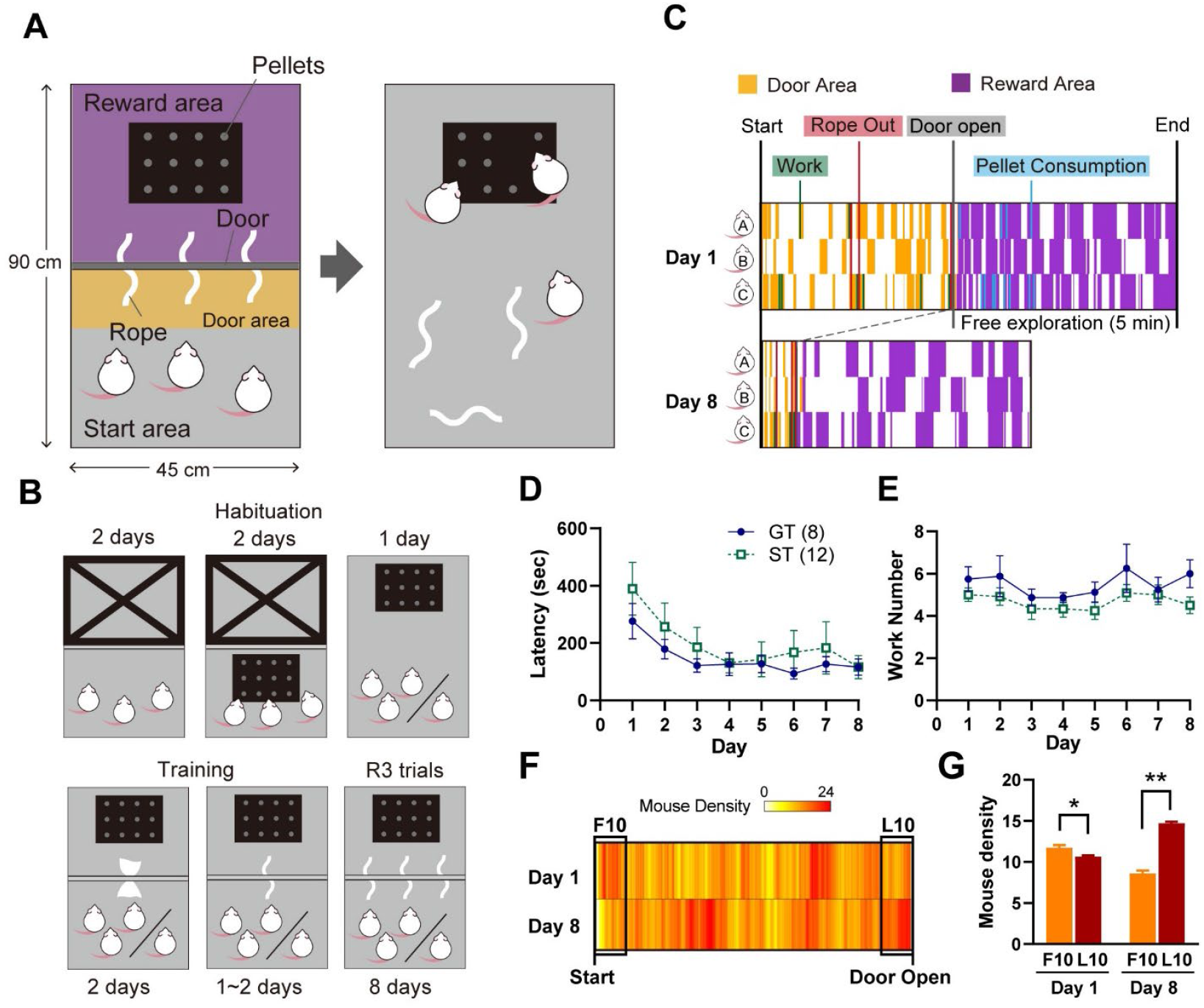
A newly developed group-based operant task for mice. **(A)** Schematic diagram of the flow of one trial of the Tsunahiki task. **(B)** Schematic diagram showing the flow of habituation and training trials and the following three-rope (R3) trials. From the 5^th^ day of habituation (top right panel), the groups were divided into two conditions: three mice indicate group-taskers (GT), and one mouse indicates single-taskers (ST). **(C)** Representative diagram showing the time course of task execution in a trial by a group of mice. Comparison between Day 1 (top) and Day 8 (bottom) of the R3 trials in the same group. The yellow (door area) and purple (reward area) indicate the time spent in each area. **(D)** Latency to all ropes in repeated R3 trials in the GT (n = 8, blue, solid circle with solid line) and ST (n = 12, green, empty square with dashed line). **(E)** Number of works in the repeated R3 trials in the GT and ST groups. **(F)** Mouse density in the door area on Day 1 (top) and Day 8 (bottom). Deep red indicates the maximum density, corresponding to a total of 24 mice in the door area. White indicates that no mice were present in the door area. The latency to door opening in each group was normalized. Black squares indicate the first (F10) and last (L10) 10 % of the trials. **(G)** Mean mouse density in the door area in the first (F10, orange) and last (L10, red) 10 % of each of Days 1 and 8. A total of 100 frames were analyzed for each time period. All data are shown as mean ± SEM, standard error of the mean. * *p* < 0.05, ** *p* < 0.001.

In the present study, we used groups of three adult male mice to examine whether mice could learn an operant task through group work experience. The results indicated that mice can quickly acquire the Tsunahiki task and spontaneously repeat the task performance, and task experience leads to an inequality of work and rewards based on the dominance hierarchy within a group. In the future, the Tsunahik task will enable us to investigate the neural basis of social relationships mediated by group work, which remains largely unclear.

## Results

### Acquisition of the Tsunahiki task in groups or individually (Experiment 1)

Adult ICR/Jcl male mice were commercially purchased (CLEA Japan) and housed in groups of three throughout the experiment. One week after being housed in groups, each group underwent habituation trials on the apparatus and reward pellets (chocolate-flavored), one trial per day, for five days (upper panels in Fig. 1B). From the 5th day of habituation, the groups were assigned to either of two conditions: group-taskers (GT; eight groups, 24 mice) or single-taskers (ST; four groups, 12 mice). Although the STs were housed in groups, they performed subsequent trials individually.

After the habituation trials, the mice were trained for rope-pulling behavior. To induce pulling behavior, a flutter paper strip (10 × 4 cm), which induces the innate behavior of gathering as a nesting material, was inserted into a hole in the door for 2 days (lower left panel in Fig. 1B). On the 3rd and/or 4th day, a polypropylene rope (15 cm length) was introduced instead of the paper strip (R1 trial) (lower middle panel in Fig. 1B). If each group or individual pulled one rope out, they proceeded to the three-rope trial (R3 trial) the next day (lower right panel in Fig. 1B). All GTs and 11 of 12 mice in the STs pulled the rope out in the R1 trial within one day, and the remaining mouse succeeded in the second R1 trial on the 4th day of training trials.

The R3 trials were conducted repeatedly for eight days (eight trials). The latency to remove all ropes gradually decreased in both conditions, indicating that the mice were able to acquire this task regardless of whether they performed it alone or in a group (Fig. 1C and D, two-way repeated measures ANOVA, Day: F_(3.047, 54.843)_ = 10.129, *p* < 0.001, Movie S1). Although there was no difference in rope-out latency between the conditions (Condition: F_(1, 18)_ = 0.376, *p* = 0.547; Interaction: F_(3.047, 54.843)_= 0.780, *p* = 0.512), the STs completed the task more efficiently, performing fewer work numbers (rope-pulling actions) than the GTs (Fig. 1E, two-way repeated measures ANOVA, Day: F_(7, 126)_ = 1.304, *p* = 0.254; Condition: F_(1,18)_ = 5.389, *p* = 0.032; Interaction: F_(7, 126)_ = 0.304, *p* = 0.951). GTs seemed to pull the rope gradually through interaction with each other and required more actions to complete the task instead of depending on a single efficient worker.

Regarding these results, one question arises: Did all mice in each group understand the task when they learned together? We found that on Day 8 in the R3 trials, mouse density in the area near the guillotine door (door area; within 15 cm of the door) increased by the time all the ropes had been removed, specifically during the last 10 % of the time before the door opened, compared to the first 10 % (Fig. 1F & G, two-way repeated measures ANOVA, Day: F_(1,99)_ = 5.936, *p* = 0.017; Time: F_(1,99)_ = 57.849, *p* < 0.001; Interaction: F_(1,99)_ = 330.815, *p* < 0.001). Multiple comparisons revealed that mice were gathered in the door area in the last 10 % compared to the first 10 % on Day 8 (*p* < 0.001), but the opposite phenomenon was observed on Day 1 (*p* = 0.02). This suggests that GTs came to understand the task flow after repeated task execution in groups.

### Group taskers can perform tasks individually

After the repeated R3 trials, the GTs underwent two individual R3 trials. Although the time until the door opened tended to increase, all mice completed the task even on the 1st day of the individual trial (Fig. 2A, t-test, R3 vs. Individual Day 1: t_(27.363)_ = 2.515, *p* = 0.054; R3 vs. Individual Day 2: t_(25.256)_ = 2.219, *p* = 0.071, paired t-test, Individual Day 1 vs. Individual Day 2: t_(23)_ = 1.890, *p* = 0.071). These results suggest that GTs chose to perform the task when faced alone.

**Fig. 2.**
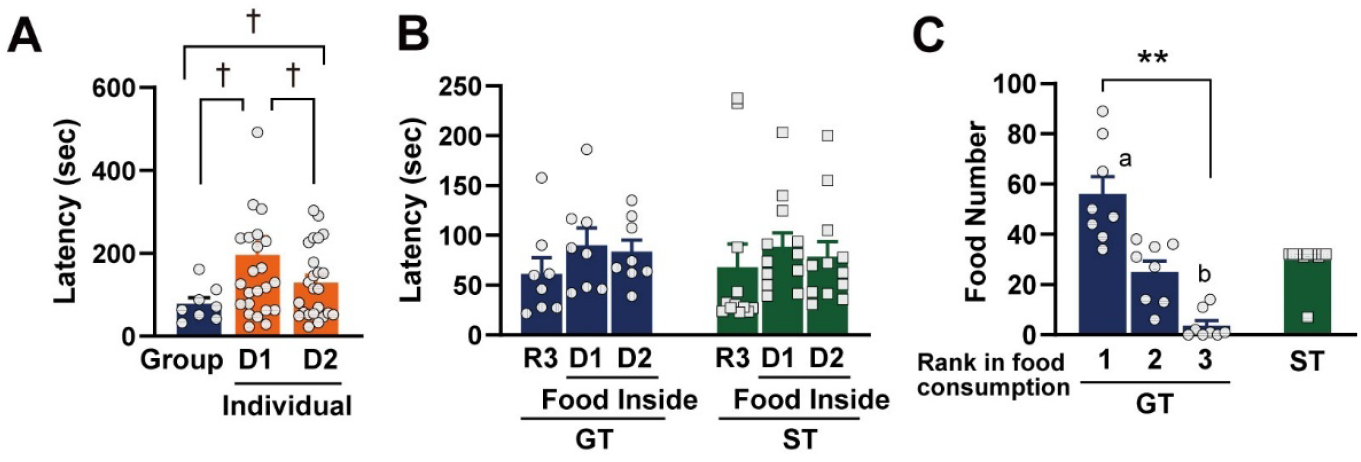
Task performance in individual trials and reward-related analyses. **(A)** Latency to all ropes out in the group (n = 8, blue) and individual (n = 24, orange) R3 trials of the GT. One data point was outside the axis limits. **(B)** Latency to all ropes in the general and food-inside R3 trials in the GT (n = 8, blue) and ST (n = 12, green). **(C)** Total food number consumed during the repeated R3 trials (total of eight trials) in each individual within a group of the GT (n = 8 each, blue) and the ST (n = 12, green). All data are shown as mean ± SEM. † *p* < 0.1, ** *p* < 0.001.

### Mice perform the task without food rewards

In this experiment, both GTs and STs ate at least one pellet in most trials, even without food restriction. To investigate whether the mice could perform the task without a food reward, we conducted R3 trials with a pellet dish in the start area (food-inside trials) for two days. Surprisingly, all GTs and STs completed the task without any food reward, although rope-out latency tended to become longer than in the R3 trial (Fig. 2B, one-way repeated-measures ANOVA, Day (GTs): F_(2, 14)_ = 2.806, *p* = 0.094; Day (STs): F_(2, 22)_ = 1.513, *p* = 0.242). These results suggest that the motivation for the mice to perform this task was entry into the reward area itself rather than eating the pellets.

### Reward distribution among group taskers

When focusing on each individual mouse, there was a large within-group variation in pellet consumption behavior in the GTs. Percentage of pellet-consuming trials, when analyzed by each mouse, showed that GTs had a significantly lower pellet-consuming rate (139/192 [72.40 %]) than STs (91/96 [94.79 %]) (X^2^ test, X^2^_(1)_ = 18.591, *p* < 0.001). Moreover, there were significant individual differences in the total number of pellets consumed over the 8-day period within a group (Fig. 2C, Kruskal-Wallis test, Rank in food consumption: H_(2)_ = 18.502, *p* < .001, rank 1 vs. rank 3: *p* < 0.001 in multiple comparison). The individual that consumed the most ate more than half of the total 96 pellets, and they ate significantly more pellets than STs (*p* < 0.001 in multiple comparisons). On the other hand, the least pellet consumers ate a significantly smaller number of pellets than STs (*p* < 0.001 in multiple comparisons). Did the mouse that monopolized the pellets work most vigorously? There was a significant positive correlation between the number of pellets consumed and the number of works in the first two days (Table 1, Spearman’s rank correlation test, *rho* = 0.365, *p* = 0.011), but it disappeared in the last two days, Days 7 and 8 (Spearman’s rank correlation test, *rho* = 0.071, *p* = 0.633). These findings suggest that social relationships within a group, such as dominance hierarchy, affect the distribution of work and reward in GTs, and that task experience may have changed their relationship.

**Table 1.** Relationship between the number of pellets consumed and work in two trials.

| Day | *rho* | *p* |
| --- | --- | --- |
| 1 and 2 | 0.365 | 0.011* |
| 3 and 4 | 0.249 | 0.088† |
| 5 and 6 | -0.092 | 0.534 |
| 7 and 8 | 0.071 | 0.633 |

† $p < 0.1$ , \* $p < 0.05$ .

### Influence of dominance hierarchy on task performance (Experiment 2)

To test the hypothesis that the dominance hierarchy within a group causes individual differences in task execution, we divided another 12 groups of ICR/Jcl adult male mice into with-rope (w/-rope) and without-rope (w/o-rope) conditions (six groups each, three mice per group). They repeated 11 days of the R3 trials for the w/-rope or control trials for the w/o-rope. Control trials were conducted using the same apparatus and procedure as the R3 trials, but without any rope pulling (Fig. 3A). To assess the social rank within each group at each timepoint, the tube test (*11*) was conducted twice, before habituation and after the end of the 11-days of repeated R3 trials (Fig. 3B).

**Fig. 3.**
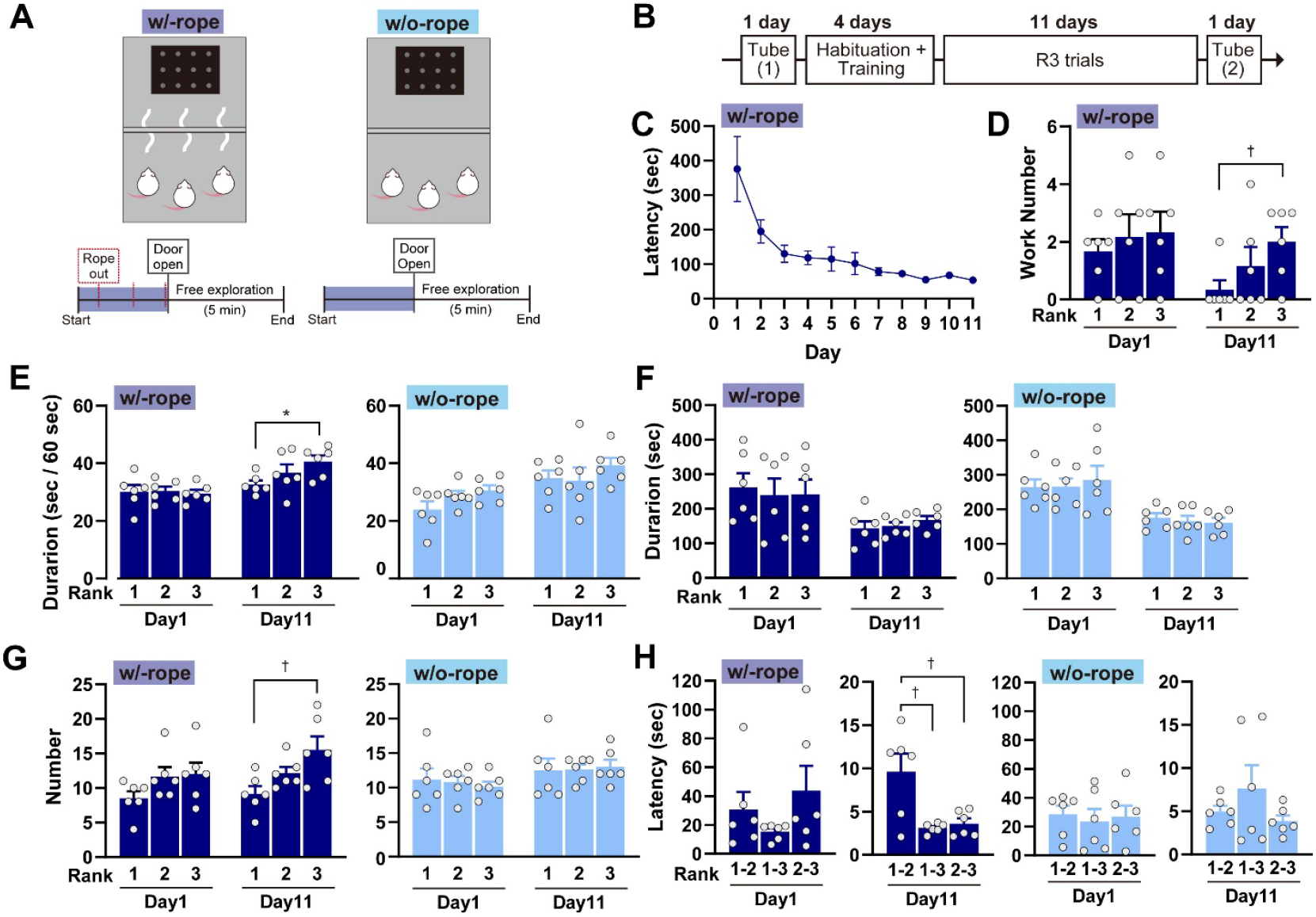
Influence of social rank on work and reward distribution. **(A)** Schematic diagram of the flow of the R3 and no-rope trials. **(B)** Schematic representation of the experimental procedures. “Tube” refers to the tube test. **(C)** Latency to all ropes out in the repeated R3 trials in the w/-rope condition (n = 6). **(D)** Number of works in the repeated R3 trials in mice of each rank (ranks 1–3) in the w/-rope condition (n = 6 for each rank) on Days 1 and 11. **(E)** Cumulative duration of staying in the door area in mice of each rank in the w/-rope (n = 6, left panel) and w/o-rope (n = 6, right panel) conditions on Days 1 and 11. **(F)** Cumulative duration of stay time in the reward area in mice of each rank in the w/-rope (n = 6, left panel) and w/o-rope (n = 6, right panel) conditions on Days 1 and 11. **(G)** Cumulative number of reward area entries in mice of each rank in the w/-rope (n = 6, left panel) and w/o-rope (n = 6, right panel) conditions on Days 1 and 11. **(H)** Latency to rank determination in the tube test in the w/-rope (n = 6, left panel) and w/o-rope (n = 6, right panel) conditions on Days 1 and 11. All data are shown as mean ± SEM. † p < 0.1, * *p* < 0.05.

### Subordinate individuals shouldered the workloads

Similar to Experiment 1, mice in the w/-rope condition showed a significant decrease in the latency to pull out all ropes across repeated trials (Fig. 3C, one-way repeated-measures ANOVA, Day: F_(10, 50)_ = 7.481, *p* < 0.001). Surprisingly, the comparison of work numbers among social ranks revealed that the individual with the lower rank within a group tended to work more than the higher rankers on Day 11, although there was no rank difference on Day 1 (Fig. 3D, Kruskal-Wallis test, Day 1: H_(2)_ = 0.621, *p* = 0.733, Day 11: H_(2)_ = 5.002, *p* = 0.082, a trend toward difference between rank 1 and rank 3 on Day 11: *p* = 0.075 in multiple comparisons). Rank 3 mice spent more time within the door area than Rank 1 mice on Day 11, but not on Day 1 (left panel in Fig. 3E, two-way repeated measures ANOVA, Day: F_(1, 15)_ = 18.594, *p* < 0.001; Rank: F_(2, 15)_ = 1.487, *p* = 0.257; Interaction: F_(2, 15)_ = 2.554, *p* = 0.111; a significant difference was observed only between ranks 1 and 3 on Day 11: *p* = 0.037 in multiple comparisons). However, the effect of task experience was not observed in the w/o-rope condition (right panel in Fig. 3E, two-way repeated measures ANOVA, Day: F_(1, 15)_ = 20.770, *p* < 0.001; Rank: F_(2, 15)_ = 1.406, *p* = 0.276; Interaction: F_(2, 15)_ = 0.827, *p* = 0.456), in which no rank difference was observed at either time point. Because most of the animals did not eat any pellets in this experiment, we used the reward area entry as an index of reward distribution. Although there was no rank difference in the duration of reward area stay even on Day 11 in both conditions ([w/-rope] left panel in Fig. 3F, two-way repeated measures ANOVA, Day: F_(1, 15)_ = 16.080, *p* = 0.001; Rank: F_(2, 15)_ = 0.042, *p* = 0.959; Interaction: F_(2, 15)_ = 0.306, *p* = 0.741; no difference among ranks, [w/o-rope] right panel in Fig. 3F, two-way repeated measures ANOVA, Day: F_(1, 15)_ = 23.196, *p* < 0.001; Rank: F_(2, 15)_ = 0.077, *p* = 0.926; Interaction: F_(2, 15)_ = 0.223, *p* = 0.803; no difference among ranks), subordinates in the w/-rope condition entered the reward area more frequently than the dominants on Day 11, but not on Day 1 (left panel in Fig. 3G, two-way repeated measures ANOVA, Day: F_(1, 15)_ = 11.200, *p* = 0.004; Rank: F_(2, 15)_ = 3.530, *p* = 0.055; Interaction: F_(2, 15)_ = 4.386, *p* = 0.032; a trend toward difference between rank 1 and rank 3 on Day 11: *p* = 0.067 in multiple comparison), suggesting that rank 3 individuals might be evicted from the reward area more frequently. In the w/o-rope condition, there was no difference in reward area entry among rank and day (right panel in Fig. 3G, two-way repeated measures ANOVA, Day: F_(1, 15)_ = 10.909, *p* = 0.005; Rank: F_(2, 15)_ = 0.014, *p* = 0.986; Interaction: F_(2, 15)_ = 0.530, *p* = 0.599).

### Task execution may promote the formation and maintenance of a dominance hierarchy

Latency to rank determination in the tube test in the 2nd test after repeated trials became dramatically shorter than that in the first test in both conditions(Fig. 3H, two-way repeated measures ANOVA, [w/-rope] Day: F_(1, 15)_ = 12.353, *p* = 0.003; Rank: F_(2, 15)_ = 1.457, *p* = 0.264; Interaction: F_(2, 15)_ = 1.401, *p* = 0.277, [w/o-rope] Day: F_(1,15)_ = 30.409, *p* < 0.001; Rank: F_(2, 15)_ = 0.028, *p* = 0.972; Interaction: F_(2, 15)_ = 0.424, *p* = 0.662). Although the decrease in latency itself seemed to be because of the effect of repeated social interaction (*12*), additional evidence suggests that task experience may have influenced hierarchical social relationships. In the w/-rope condition, the rank tended to be determined faster between ranks 1-3 (*p* = 0.077) and 2-3 (*p* = 0.077) than between ranks 1-2 in multiple comparisons, showing rank disparities parallel to the concentration of work in rank 3 individuals. These phenomena were not observed under the w/o-rope condition. Moreover, a longer latency to rank determination between two mice, which can be interpreted as a smaller disparity, may indicate the proximity of the relationship. Focusing on each condition, if the latency to rank determination between ranks 2-3 was longer, these two animals stayed in the reward area longer in the w/-rope condition (Table 2, Spearman’s rank correlation test, *rho* = 0.886, *p* = 0.019), although the w/o-rope condition showed no positive correlation (Spearman’s rank correlation test, *rho* = -0.771, *p* = 0.072). These correlations between the rank determination latency and reward-distributing behaviors were observed on Day 11, but not on Day 1, even in the w/-rope condition (Spearman’s rank correlation test, [w/-rope] *rho* = -0.314, *p* = 0.544, [w/o-rope] *rho* = - 0.029, *p* = 0.957). These results suggest that task experience promotes the establishment of relationships within a group and enhances the influence of the dominance hierarchy on the distribution of work and rewards.

**Table 2.** Relationship between rank determination latency and staying time together in the reward area.

| Rank | w/-rope |  |  |  | w/o-rope |  |  |  |
| --- | --- | --- | --- | --- | --- | --- | --- | --- |
|  | Day 1 |  | Day 11 |  | Day 1 |  | Day 11 |  |
|  | *rho* | *p* | *rho* | *p* | *rho* | *p* | *rho* | *p* |
| 1-2 | 0.143 | 0.787 | -0.029 | 0.957 | 0.543 | 0.266 | 0.143 | 0.787 |
| 1-3 | 0.086 | 0.872 | -0.486 | 0.329 | 0.371 | 0.468 | -0.029 | 0.957 |
| 2-3 | -0.314 | 0.544 | 0.886 | 0.019* | -0.029 | 0.957 | -0.771 | 0.072† |

† $p < 0.1$ , \* $p < 0.05$ .

## Discussion

In the present study, we established the Tsunahiki task, a group-based operant task for mice, in a laboratory setting. We demonstrated that adult male mice could perform the task either in groups or individually. When mice perform tasks in a group, they can freely distribute their workloads and rewards. Moreover, each individual can choose whether to contribute to task completion in a partially independent manner from the distribution of the reward. Thus, the behavior of each mouse can reflect the social context and relationships within a group. We found that task performance was influenced by the dominance hierarchy; subordinates spent more time in front of the door to shoulder a greater workload. Therefore, the Tsunahiki task enables us to investigate the influence of hierarchical social relationships on the unequal distribution of work and rewards in laboratory rodents.

Cooperation and collaboration are essential for the survival of social species. For example, collaborative hunting leads to successful foraging in killer whales, where they show division into “striker” and “helper” (*13*). The Tsunahiki task is groundbreaking because it enables the reproduction of collaborative scenarios involving three or more individuals in the laboratory. In the present study, the group did not show an equal distribution of work and rewards after the repetition of eight or eleven trials, which is in line with the role division in previous studies. We found that subordinates, especially the third, the lowest rankers, were primarily fixed to the role of “worker” in the Tsunahiki task. To observe collaborative behavior in mice, another task was performed in which a group of five to ten mice seized food from a predator-like spider-shaped robot (*14*). In that task too, mice distribute their work related to hierarchical social relationships within a group. Contrary to our findings, middle-rankers within groups worked most actively. This may be because the lowest rankers lacked the coordinated motor skills required for task execution. It should be noted that, similar to hunting in killer whales, foraging against the spider robot is fear-evoking and requires sufficient motor coordination. Our Tsunahiki task does not require high levels of motor coordination and is designed to observe collaborative social interactions in more peaceful and non-fear-inducing situations, such as group work in university classrooms. In contrast to the collaborative behavior in this study, cooperative behavior that requires precise timing synchronization among multiple individuals has been vigorously investigated in previous studies using a pair of animals. In the loose-string task, two animals are required to pull a string simultaneously to acquire a reward. This task was first established in chimpanzees *(Pan troglodytes)* (*9*), and it has been demonstrated that a wide variety of animal species, including macaque monkeys (*15, 16*), wolves (*17*), and porcupines (*18*), can perform the task. Although mice seemed unable to perform the loose-string task, a pair of mice can perform a same-typed cooperative task in which they are required to stay simultaneously beneath the straws for a liquid reward (*10*). Unlike these experimental paradigms, the Tsunahiki task requires each individual to contribute to the progression of the task without precise synchronization while observing other animals’ behavior. Similar to human group work research in modern society, the Tsunahiki task requires collaboration where the entire group shares goals and rewards, yet not everyone acts with a strict timing agreement. Reproduction of the specific situation, which is comparable to human group-work studies, in mice, the Tsunahiki have paved the way toward elucidating the neural basis of context-dependent control of social behavior.

The Tsunahiki task cannot exclude the possibility that some individuals will lose their opportunity for task acquisition because the task can be completed quickly by a particular individual in all trials. However, this was not the case in the present study.

Throughout Experiments 1 and 2, all mice worked at least two times over eight (Experiment 1) or eleven (Experiment 2) trials. Collectively with the previous findings indicating that mice are capable of observational learning (*19*) and with our observation that mice began gathering in front of the door right before the task completion after the task experiences, it can be suggested that the group taskers could acquire the task sequence throughout repetition. Thus, mice that did not show active engagement in the task may have decided not to engage rather than not being able to perform the task. However, it should be noted that when the task was performed individually by the GTs, there were large individual differences in the latency to pull out all ropes, although all members of the group taskers completed the task from the first trial. The latency in the second individual R3 trial was significantly and negatively correlated with the cumulative work number in the previous eight trials performed in groups (Fig. S1); that is, the mouse that experienced more work in its group seemed to complete the task more quickly when it performed the task individually. These results suggest that experience-dependent skill acquisition through hard work in groups also exists in mice, similar to natural human settings.

We further demonstrated that sharing work and rewards in the repeated Tsunahiki task facilitated rank discrepancy within a group. Comparison between with and without rope conditions revealed that experience of the repeated Tsunhaiki task trials promoted the establishment of a dominance hierarchy in a group. Hierarchical social relationships in male mice are believed to be stabilized within a few days (*20, 21*) based on observations from previous studies conducted in a semi-natural environment. Although it has also been reported that social rank becomes stable, that is, the same results are obtained from the repeated tube test in four consecutive days, within two weeks in a general laboratory housing environment (*22*), another study reported that not all cages maintain stable dominance hierarchies over long periods of time in the laboratory setting (*23*). In the present study, we initiated habituation trials 10 days after purchase. Collectively with the fact that mice were kept in shoebox cages in a general laboratory environment and that the w/o-rope condition showed frequent rank changes, it is likely that the Tsunahiki task promoted rank consolidation during the period when the hierarchy was unstable. This finding is in line with previous reports in humans (*3, 4*)

Rank-dependent differences in reward acquisition, with higher rankers acquiring more, were observed in this study. On the other hand, lower rankers shouldered more work without commensurate rewards. Here, one question arises: how was this inequality established? Although we initially hypothesized that the dominant animals drove out the third-ranked animal, we rarely observed chasing behavior, and the subordinates seemed to leave the reward area spontaneously. Moreover, we rarely observed aggression, even after repeated trials. It is usual that dominant individuals in social species, including mice, acquire larger resources, including food and mates (*5, 24, 25*), and this does not always occur through dominant individuals forcibly subjugating subordinates. For example, when two monkeys were faced with one food resource, the subordinate monkey spontaneously gave up eating after once social rank was determined (*26*). Similarly, third-ranked mice also seemed to spontaneously give up continuously staying in the reward area in the present study. The relationship between rank-determination latency in the tube test and staying time in the reward area in rank 2 and 3 mice, that is, a smaller rank discrepancy was related to a longer staying time together, may provide an explanation. That is, the difficulty of staying together with higher-ranking individuals with greater rank disparity may have driven them out of the reward area.

We placed chocolate-flavored pellets in the reward area without food restrictions. Nevertheless, even for the group that consumed all the pellets, entrance to the reward area seemed to be the primary reward in this task. This is evidenced by the fact that all groups opened the door when the pellet dish was placed in the start area. Even after the repetition of the same task trials, all individuals invariably entered the reward area once the door opened, suggesting that they did not become bored after at least 15 repetitions. However, since only one trial per day was conducted this time, it remains unclear whether repeating the trials within a single day would cause the mice to stop performing the task. Although many aspects of why mice perform this task remain unclear, it is possible that the task itself is enjoyable, similar to the hide-and-seek tasks for rats (*27*). Nevertheless, considering that the correlation between pellet consumption and work number was only observed during the initial phase of the task in Experiment 1 and that work performance shifted to subordinate individuals as the task was repeated, it is highly likely that work itself did not remain rewarding indefinitely and ultimately became a means of gaining access to the reward area.

Taken together, the Tsunahiki task, a group-based operant task developed in this study, provides a novel behavioral paradigm for observing group work in laboratory rodents. This enabled us to compare individual and group conditions and investigate the effects of working together on rodents. In this study, we used adult male mice for the experiments. It is known that females also establish their hierarchy, and the form of hierarchy is not completely consistent with that of males (*28*). The estrous cycle may affect task performance related to hierarchical social relationships. We will conduct similar studies on females to clarify how they perform tasks and distribute work and rewards in the future. The Tsunahiki task does not require food or water restriction or punishment. Thus, this task is not a model for the context of cooperation for survival, such as war, but for a more peaceful context of group work where the majority of social issues in modern society arise. The Tsunahiki task will contribute to elucidating the neural mechanisms related to these issues and lead to the development of biological interventions.

## Materials and Methods

### Animals and housing conditions

Adult male ICR/Jcl mice were purchased from a commercial breeder (CLEA Japan Inc., Japan). On the day of arrival, the mice were assigned to groups of three, each consisting of individuals who had been transported in the same cage and were similar in weight. Behavioral experiments were initiated one week after arrival. Mice were housed under standard housing conditions at a room temperature of 23±2 °C. Food and water were available *ad libitum*. All behavioral tests were conducted in the dark phase of a 12-h light–dark cycle, and the mice were moved to the experimental room more than one hour before the test started. All behavioral tests were conducted under dim white light (10 ∼ 15 lux). All experiments were conducted in accordance with the NIH guidelines. All efforts were made to minimize the number of animals used and their suffering.

In experiment 1, 36 mice (12 groups) were 9 weeks old and weighed 34.45±1.45 g at the start of the experiment. The weight difference within a group was within 3.09 g at the start. This experiment was conducted at the National Institute of Advanced Industrial

Science and Technology (AIST). Animals were kept in regular plastic housing cages (18 × 29 × 12 cm) with flatter paper bedding and general mouse food. The light was turned off at 3 pm, and behavioral tests started at 5 pm. All experiments were approved by the Animal Care and Use Committee of AIST.

In experiment 2, 36 mice (12 groups) were 10 weeks old at the start of the experiment and weighed 38.02±2.99 g at the start. Weight difference within a group was within 1.37 g at start. This experiment was conducted in the University of Tsukuba.

Animals were kept in plastic regular housing cages (18.5 × 28.7 × 12.5 cm) with corncob bedding and general mouse food. Short transparent acrylic tube (inner diameter 3 cm and 10 cm length) was introduced in each home cage for habituation to the tube used in tube test from the day of arrival. The short tubes were removed before the first habituation trial. The light was turned off at noon, and behavioral tests were started at 2 pm. All experiments were approved by the Animal Care and Use Committee of the University of Tsukuba.

### The Tsunahiki task

#### Apparatus

The task was conducted in a gray opaque acrylic open-field box (90 × 90 × 40 cm, LE800SC, Panlab). The box was divided in half by acrylic panels of the same color, with one half used as the experimental field. The field was further divided in half by a transparent acrylic guillotine door in the start and reward areas. The door was perforated, allowing the mice to sniff beyond it. Six holes with a diameter of 1 cm were placed at a height of 3 cm from the floor and lined up at roughly equal intervals. A paper strip or rope was inserted into the holes for the task (Fig. S2).

In the reward area, a black acrylic plate (21 × 22 cm) with chocolate-flavored sucrose pellets (Bio-Serv, 20 mg) was placed. It should be noted that the plate was placed in the start area for the food-inside trials. The dish had 12 round indentations, each 2 cm in diameter. A total of 12 pellets (one pellet per indentation) were used in a trial with three mice. When a trial was performed with a single mouse, four pellets were placed in the center row.

#### Procedures

At the beginning of the task, the mice were released from a black opaque plastic start box (14 × 14 × 12 cm) placed at the edge of the start area. They pulled on a paper strip (paper trial, Kimwipe(R) strip in 10 × 4 cm was used), a rope (R1 trial), or all three ropes (R3 trial) that were stuck through the hole(s) in the door. When they pulled out all the ropes from the door hole(s) within 20 minutes, the guillotine door was opened, and the mice could explore the reward area, where reward pellets were placed, except for the food-inside trials, for 5 minutes. All animals could enter the reward area, regardless of which individual executed the task. Similarly, the pellet acquisition was independent of individual task performance.

If the rope accidentally fell into the reward area before being pulled out to the start area, the experimenter quickly returned the rope to its original position. If the entire rope was pulled out into the start area, the trial was considered to have been completed, and the door was opened even if the mice did not appear to actively pull out the rope with their forelimbs or mouth.

The w/o-rope trials in Experiment 2 were conducted using the same procedure as the R3 trial but without any ropes. The door was opened by an experimenter in a time-dependent manner (see the *Experimental procedures* for *Experiment 2*). In the food-inside trials in Experiment 1, the pellet dish was placed at the center of the start area.

After the end of each trial, the field and start box were cleaned with 70 % ethanol and then wiped with water to remove any ethanol odor. For each trial, a new piece of paper or a new set of ropes was used.

### Behavioral analysis

All behavioral tests were recorded using digital video cameras placed above the experimental field, and all video recordings were scored by an experimenter using a digital event recorder program (Recordia 1.0b, O’Hara). Entry to the door and reward area and food consumption were observed and recorded individually. The number of “work,” which was defined as an active rope-moving behavior using forelimbs and mouth, was also counted for each individual. The latency to each rope falling out into the start area was recorded for each trial, even if it was not induced by work.

### Tube test

Tube tests were conducted to assess the rank of each animal within each group, using procedures based on previous studies (*5, 12*). A transparent acrylic tube (inner diameter 3 cm and 30 cm length) was placed in the same experimental field as the rope task and was used for testing. Mice were trained for two days and then underwent test trials. Because the test was conducted in a round-robin tournament within each group, it was performed three times per group in a single session. Each mouse was trained to run through the tube for two consecutive days before the first test. Each mouse was placed in a tube and allowed to move back and forth within the tube four times per day. If the mouse attempted to exit the tube by backing up, the experimenter gently pushed it back into the tube. Training was conducted on each day of testing. The mice made two round trips through the tube in each practice trial conducted immediately before the testing trial. In the testing trial, two mice were placed at each end of the tube, and they were released to start when the entire body of both mice entered the tube. When all limbs of one mouse emerged from the tube, the game was judged to be over. The individual that emerged from the tube was a loser (subordinate), and the individual that remained in the tube was a winner (dominant). If no mouse was ejected within two minutes, the trial was considered a tie; however, no trial resulted in a tie in this experiment.

### Experimental procedures

#### Experiment 1

One week after arrival, mice were habituated in each group to the experimental apparatus for 20 minutes per day for a total of four days, one trial per day. The guillotine door was replaced with a gray opaque acrylic wall, and the mice were allowed to freely explore the start area of the open field. The pellet dish with 12 pellets per trial was introduced in the last two trials during habituation. On the 5th day of the habituation trial, the guillotine door was opened, and a pellet dish was placed in the reward area. In this trial, the mice were allowed to freely explore both the start and reward areas for 15 minutes. After this trial, the groups were assigned to one of two conditions: group-taskers (GT; eight groups, 24 mice) or single-taskers (ST; four groups, 12 mice). Mice in the ST condition underwent trials individually, although they were kept in groups.

The mice were then trained for the rope task after the habituation trials. Exploiting the mouse’s habit of pulling in fluttering objects, one paper strip was inserted into a hole in the middle of the guillotine door. Regardless of the results of the two paper trials, all mice proceeded to a trial in which the paper strip was replaced with a single 15-cm rope (R1 trial). All GT and ST succeeded in pulling out the rope within two R1 trials and proceeded to R3 trials. After the end of eight days of R3 trials (one trial per day), all mice individually performed the R3 trials (individual trials), regardless of the condition, for two consecutive days. Food-inside R3 trials were conducted for two consecutive days. One general (in a group for GT, individually for ST) R3 trial was conducted right before day 1 of the individual and food-inside trials as a control trial. The trials were not conducted consecutively; after the consecutive trials up to 5 trials, a break of two days was employed.

#### Experiment 2

In this experiment, mice were first trained for the tube test for two days and then assessed for their ranks using the tube test in a round-robin tournament within the group. They were then placed in each group in the experimental apparatus for 10 minutes per day for 2 days, with the guillotine door closed. In the third habituation trial, the door was opened. Twelve pellets were placed on the pellet dish in each trial, and the dish was placed in the start area in the initial two trials, but it was placed in the reward area in the third.

To examine the effect of task execution, the groups were assigned to one of two conditions: with-rope (w/-rope) condition or without-rope (w/o-rope) control condition (six groups each, three mice per group). The mice assigned to the w/-rope condition performed the Tsunahiki task as in Experiment 1, whereas the mice assigned to the w/o-rope condition performed a control task in which the door was opened by an experimenter without any rope pulling (no-rope trials). To control the latency to the door opening, each group of the w/o-rope condition was paired with each group of the w/-rope condition. The door for the w/o-rope condition was opened at the same latency as the paired group in the w/-rope condition on each experimental day.

The groups assigned to the w/-rope condition were trained for the Tsunahiki task using a single 15-cm rope, which was spread, fluttered, and inserted into a hole in the middle of the guillotine door (R1 trial). All groups succeeded in pulling out the ropes in a single trial and then proceeded to R3 trials. The groups in the w/-rope condition then repeated 11 days of the R3 trials, whereas the groups in the w/o-rope condition performed 11 days of no-rope trials. After 11 days of R3 or no-rope trials (one trial per day), all groups were tested again using the tube test to assess their ranks within each group. The trials were not conducted consecutively; after the consecutive trials up to 5 trials, a break of two to three days was employed.

#### Statistical Analysis

Statistical analyses were performed using IBM SPSS Statistics (version 29.0.0.0) and R (version 4.1.0)(*29*). In experiment 1, the latency to all ropes out, number of work in a trial, and mouse density in the door area were analyzed using two-way repeated measures ANOVA. For the analysis including individual trials, the latency to all ropes out was analyzed by either an unpaired t-test (comparison between the R3 and individual trials) or a paired t-test (comparison between individual trials). For food-inside trials, the latency to all ropes out was analyzed using one-way repeated-measures ANOVA for each group. The number of pellets consumed was analyzed using the Kruskal-Wallis test (within a group) or multiple comparisons using the U-test (versus ST individuals). The relationship between the number of pellets consumed and work number was analyzed using Spearman’s rank correlation. The X^2^ test was used to compare the rate of pellet-consuming trials. In experiment 2, the latency to all ropes out was analyzed using one-way repeated measures ANOVA, the number of work was analyzed using the Kruskal-Wallis test for each day, staying duration in the reward and door areas, number of the reward area entry, and the latency to the rank determination in the tube test were analyzed using two-way repeated measures ANOVA for each group. The relationship between rank-determination latency and co-stay duration was analyzed using Spearman’s rank correlation. Greenhouse-Geiser correction was used for the degree of freedom in repeated measures ANOVA, if necessary. Multiple comparisons were conducted using Holm’s p-value adjustment. Differences were considered significant at *p* < 0.05 (*), *p* < 0.01 (**), and *p* < 0.001 (***).

## Acknowledgments

We thank Dr. S. Yamamoto, Dr. S. Ogawa, Dr. S. Yamane, Dr. Y. Watanabe, Dr. I. Takashima, S. Koshimizu, and K. Ohara for valuable discussions, S. Inudsuka and K. Kodama for support for technical assistance, Dr. S. Sagoshi provided an illustration of experimental apparatus in the supplemental figure. We also thank A. Sagehashi for secretarial assistance. English-language proofreading was conducted using Paperpal AI by Editage.

## Author contributions

Conceptualization: MN, RI

Methodology: MN, RI, TS

Investigation: MN, RI, SD, TM, KH, YK, TS

Project administration: MN, TS

Writing—original draft: MN

Writing—review & editing: MN, RI, TS

## Data and materials availability

All data in the main text or the Supplementary Materials are available from the corresponding authors upon reasonable request

