## Supplementary material for "Tsunahiki task: A newly developed group-based operant task for mice": Figure S

Mariko Nakata *et al.*

*Corresponding author. Email:



**This PDF file includes:**

Figs. S1 to S2
Caption for Movie S1

**Other Supplementary Materials for this manuscript include the following:**

Movie S1

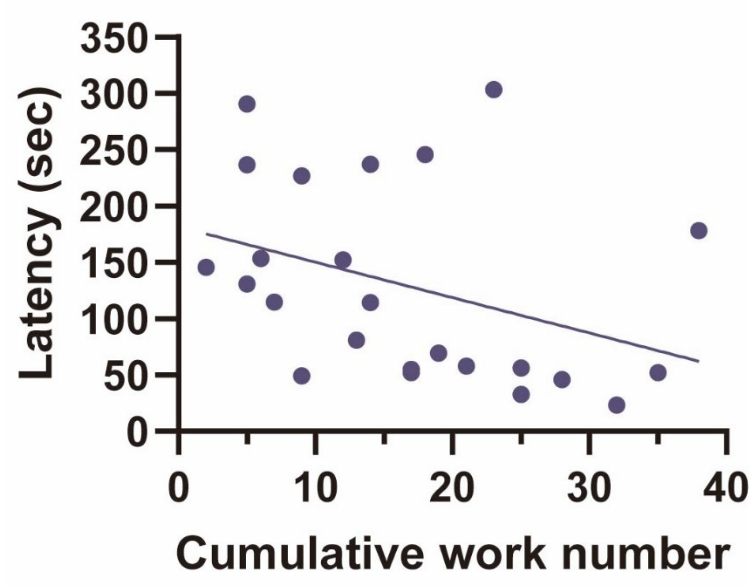


**Fig. S1.**

Relationship between the latency to all ropes out on Day 2 of the individual trials and the cumulative work number in the repeated R3 trials (a total of eight trials). Each point indicates a mouse in the GT condition (n = 24). The latency to all ropes out on Day 2 of the individual trials showed a negative and moderate correlation (*rho* = -0.443, *p* = 0.030*) with the cumulative work number in the repeated R3 trials**.**

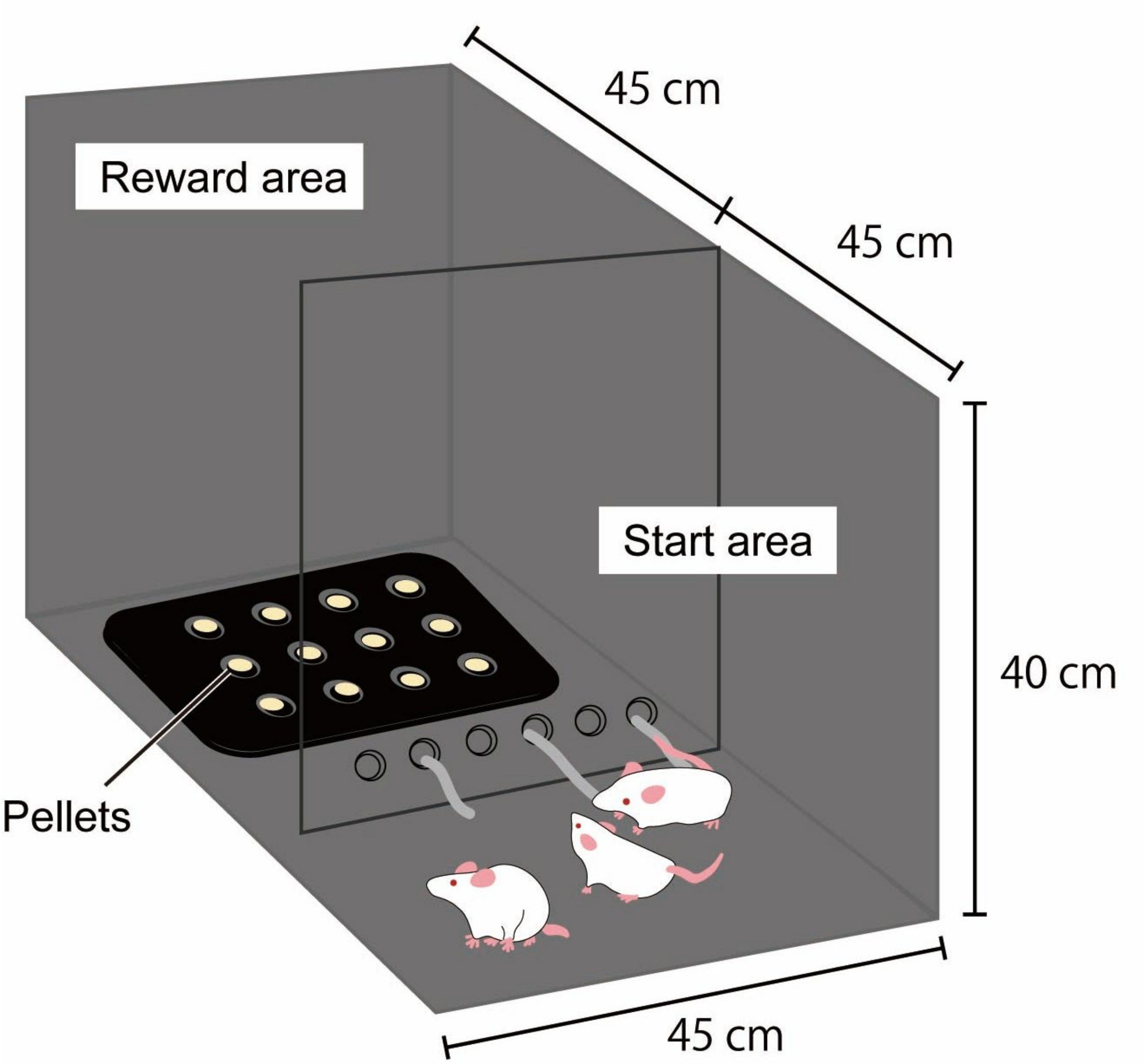


**Fig. S2.**
Schematic representation of the experimental apparatus.

**Captions for Supplementary Movie 1.**

Movie S1. Behavioral events in one R3 trial. The behavioral and event labels are presented at the bottom. This movie shows the behavioral events throughout the entire period until the door opens and from the door opens until the mice finished consuming all the pellets. The video speed was increased to 2x, as indicated at the bottom right.
